# Identification of promising oilseed rape genotypes for the tropical regions of Iran using multivariate analysis

**DOI:** 10.1101/2021.02.23.431199

**Authors:** Behnam Bakhshi, Hassan Amiri Oghan, Bahram Alizadeh, Valiollah Rameeh, Kamal Payghamzadeh, Davood Kiani, Mohammad Rabiee, Abbas Rezaizad, Gholamhossein Shiresmaeili, Alireza Dalili, Shahriar Kia

## Abstract

producing new adapted oilseed rape cultivars among the available resources of rapeseed would be a valuable method to increase the cultivar diversity in the tropical regions. Low adaptable and high yield cultivars resources of oilseed rapes are now available in the tropical regions of Iran. The current study aimed to identify new high yield and adaptable genotypes adaptable across various tropical regions. To this end, 20 new genotypes and a check variety (Dalgan) were cultivated in the tropical regions of Iran based on a randomized complete block design (RCBD) with three replications during the 2019 to 2020 cropping season. The experimental sites are composed of five locations, including Gorgan, Sari, Rasht, Borazjan and Zabol. During the growth season, several phenological and quantitative traits were recorded. Combined ANOVA revealed significant genotype by environment interaction for all studied quantitative traits. Days to start flowering and days to end flowering showed the highest heritability. Correlation analysis showed a significant positive relationship between yield and flowering period, the number of sub-branches and also the number of pods per plant, but a negative and significant correlation with the days to maturity. Path analysis showed that the days to maturity had the most negative direct effect on yield and the days to start flowering, while the number of sub-branches had the most positive direct effect on yield. Canonical correlation showed that yield is correlated positively with phenological traits. The principal component analysis showed that the two first components covered 68.07% of all data variations which 12 genotypes were correlated with these two components. Cluster analysis categorized evaluated genotypes into three main groups. Finally, eight genotypes (from class 2 of the cluster) were selected in the current study, which had high yield and adaptability in the tropical regions of Iran.

## Introduction

Rapeseed (*Brassica* species) has been cultivated to produce oil for thousands of years(Canada, 2013). Global demand for oilseed rape has steadily increased in the last 60 years, and oilseed rape has now become the second largest oilseed crop after soybean, with more than 22.7 million tons of oilseed rape annual production(FAO, 2016).

Ongoing climate change has been introduced as the main limiting factor in oilseed rape production in many part of the world (Lobell & Gourdji, 2012). The global concern in oil production is mainly focused on increasing drought stress, heat stress, or a combination of both stresses(IPCC, 2007), which could lead to oilseed rape yield reduction(Wu & Ma, 2018). As an example, 3-4°C temperature incensement leads to 15-35% of yield loss in the Middle East (Ortiz *et al.*, 2008). Therefore, one of the principal aims of crop breeding is the genetic improvement and climate-adapted cultivar development (Fischer & Edmeades, 2010). For instance, several early flowered cultivars with sufficient vegetative biomass to support seed filling have been developed in Western Australia under drought stress (Thurling, 1991). Manipulating the number of days to flowering could lead us to achieve adapted cultivars across the different environmental (Diepenbrock, 2000).

Iran is one of the mail oilseed rape producers in the Middle East, which is ranked as 27^th^ country in terms of the harvested area of canola in the world(FAO, 2018). However, oilseed rape cultivation in Iran is facing heat and drought stress, especially in tropical regions, while *Brassica* species are adapted to high rainfall areas. Therefore, producing newly adapted genotypes among the available resources of rapeseed would be a valuable method to increase the cultivar diversity in the tropical regions. Several studies have been down to identify adopted cultivars of oilseed rape. Aghaei(Agahi *et al.*, 2020) examined 22 genotypes of oilseed rape at five experimental sites and reported that Hyola 401 had the highest yield and adaptability. Likewise, eleven new oilseed rapes were evaluated in the seven different environments and the RBN-04722 genotype was identified as the most adaptable genotype(Tahira & Amjad, 2013).

Oilseed rape genotypes showed various yields across different environments because of their interaction with the environment. The problem is that yield is a complex characteristic that is regulated by many genes and highly affected by environmental factors. The different experimental environments should be considered to identify genotypes with the highest yield and the lowest fluctuations in the different environments(Agahi *et al.*, 2020). Evaluation of yield in the different environments could help us select high yield and stable genotypes with high accuracy (Roy, 2000). Recently 17 new oilseed rape lines were examined in the different experimental sites of the south tropical regions of Iran. Finally, three high yield genotypes adapted with the tropical regions of Iran were identified (Amiri Oghan *et al.*, 2019). Therefore, to introduce new promising genotypes, various new identified lines should evaluate across different environments. In the current study, 20 genotypes with one check genotype are evaluated in the five experimental locations of the tropical regions, including Gorgan, Sari, Rasht, Borazjan and Zabol, to detect high yield and adaptable genotypes.

## Material and Methods

In the present study, 20 new genotypes and a check variety (Dalgan) were cultivated in the tropical regions of Iran based on a randomized complete block design (RCBD) with three replications during the 2019 to 2020 cropping season. The experimental sites are composed of five locations, including organ, Sari, Rasht, Borazjanand Zabol, which are considered tropical regions of Iran (Table 1). The physical and chemical characteristics of the soil at the experimental field were examined before and during the experiment. The evaluated genotypes were composed of 20 pure line genotypes, including G1-G20, which were achieved by pedigree breeding projects in the seed and plant improvement institute of Iran (SPII). The pedigree of all genotypes is presented in table 2. Dalgan variety, which is a spring type of oilseed rape cultivar, was utilized as check variety.

**Table 1.** Geographical characteristics of the experimental locations.

| Locations | Altitude (m) | Longitude | Latitude | Rainfall (mm) |
| --- | --- | --- | --- | --- |
| Gorgan | 5.5 | 54.20 | 36.55 | 400-500 |
| Sari | 29 | 53.10 | 36.41 | 650 |
| Rasht | 10 | 49.27 | 37.27 | 1359 |
| Zabol | 489 | 61.32 | 31.5 | 60 |
| Borazjan | 80 | 51.17 | 29.16 | 283 |

**Table 2.**
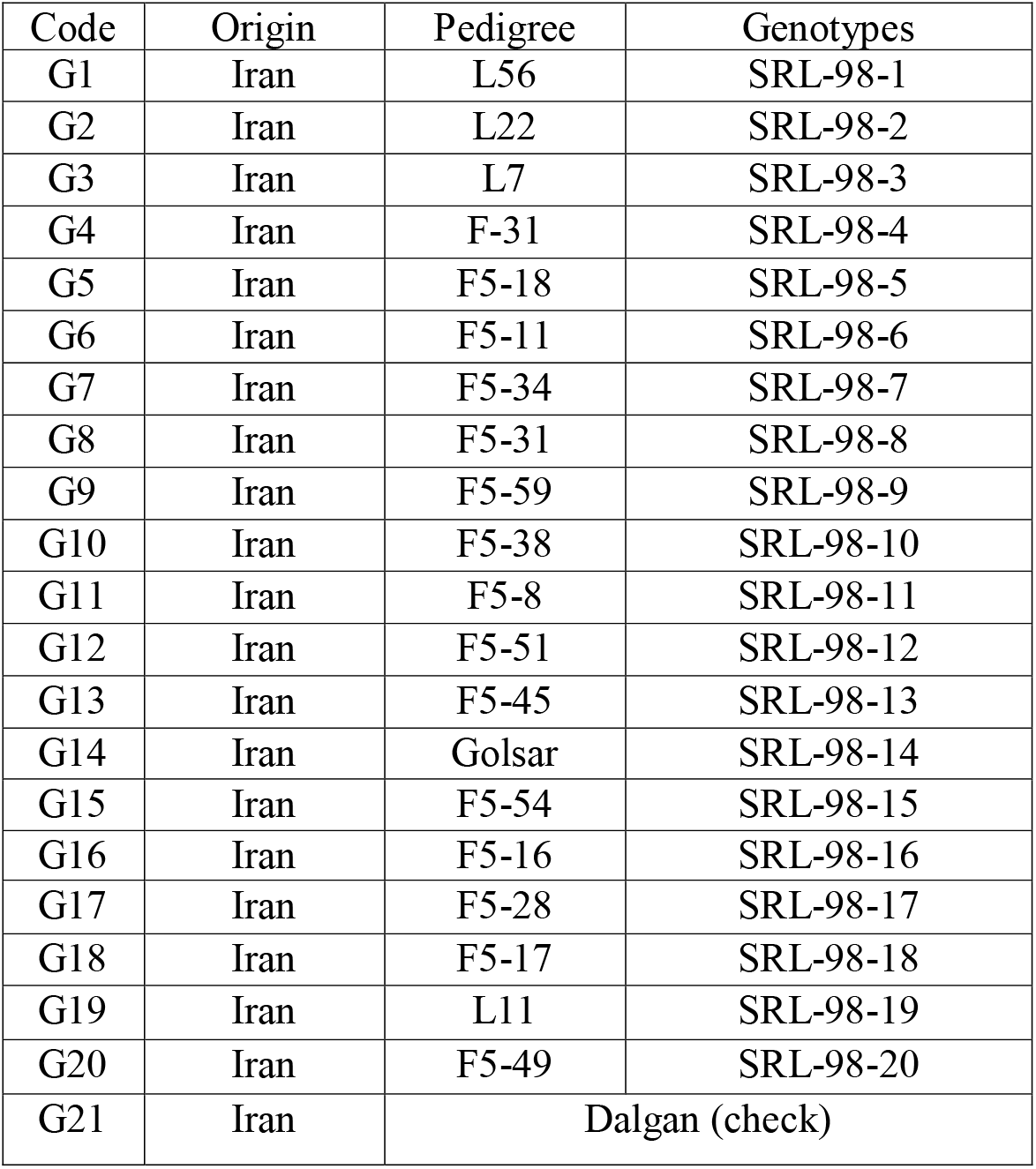
Pedigree of the pure lines evaluated in five experimental locations.

| Code | Origin | Pedigree | Genotypes |
| --- | --- | --- | --- |
| G1 | Iran | L56 | SRL-98-1 |
| G2 | Iran | L22 | SRL-98-2 |
| G3 | Iran | L7 | SRL-98-3 |
| G4 | Iran | F-31 | SRL-98-4 |
| G5 | Iran | F5-18 | SRL-98-5 |
| G6 | Iran | F5-11 | SRL-98-6 |
| G7 | Iran | F5-34 | SRL-98-7 |
| G8 | Iran | F5-31 | SRL-98-8 |
| G9 | Iran | F5-59 | SRL-98-9 |
| G10 | Iran | F5-38 | SRL-98-10 |
| G11 | Iran | F5-8 | SRL-98-11 |
| G12 | Iran | F5-51 | SRL-98-12 |
| G13 | Iran | F5-45 | SRL-98-13 |
| G14 | Iran | Golsar | SRL-98-14 |
| G15 | Iran | F5-54 | SRL-98-15 |
| G16 | Iran | F5-16 | SRL-98-16 |
| G17 | Iran | F5-28 | SRL-98-17 |
| G18 | Iran | F5-17 | SRL-98-18 |
| G19 | Iran | L11 | SRL-98-19 |
| G20 | Iran | F5-49 | SRL-98-20 |
| G21 | Iran | Dalgan (check) |  |

Each experimental plot was designed with four lines of 5 meters at a distance of 30 cm and a row spacing of 30 cm. Plant density was about 60 plants per m^2^. Fertilization was performed according to local customs after land preparation. During the five phases, irrigation was carried out, including sowing, rosette, flower initiation, initiation of pods, and development of grain. During the growth season, several phenological traits including days to start flowering (DSF), days to end flowering (DEF), flowering period (FP), days to maturity (DM) and also several yield component traits including one-thousand seed weight (TSW), number of sub-branches (NSB), number of grain per pods (NGP), number of pods per plant (NPP) and yield were recorded.

In the current research, all investigated quantitative traits were analyzed using combined ANOVA for the five experimental locations. Correlation analysis has been conducted using the Pearson method. In order to investigate the relationship between phenological traits and yield components, the canonical correlation has been used. Additionally, Path analysis was utilized to find the direct and indirect effect of quantitative traits on yield. Furthermore, cluster analysis using the WARD method and also principal component analysis was utilized to classify genotypes. In order to analyze the data, IBM SPSS Statistics ver. 24 and XLSTAT ver. 2019.2.2 computer programs were used.

## Results and Discussion

Combined ANOVA revealed significant genotype by environment interaction for all studied quantitative traits as well as the environments. So, the genotypes showed different performances across the environments. Genotypes were also significant for all their phenological traits and yield at 1% level and one-thousand seed weight at 5%. Genotypes were not significant for other yield component traits, including the number of sub-branches, the number of grain in pods and the number of pods per plant. Therefore, the studied genotypes are more diverse in their phenological traits; however, when genotypes interacted by the environment, all yield components were also significant, which shows the strong influence of the environment in the yield component of the evaluated genotypes. The low and high coefficient of variation was also observed for phenological and yield components, respectively. The high value of the yield coefficient of variation confirmed the effects of the environment on yield components. On the other hand, while all the test environments were located in the tropical regions, a lower coefficient of variation was observed for phenological traits (Table 3).

**Table 3:**
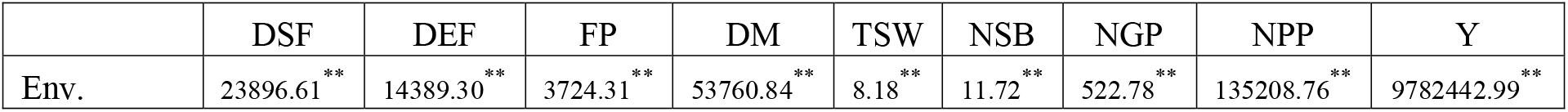
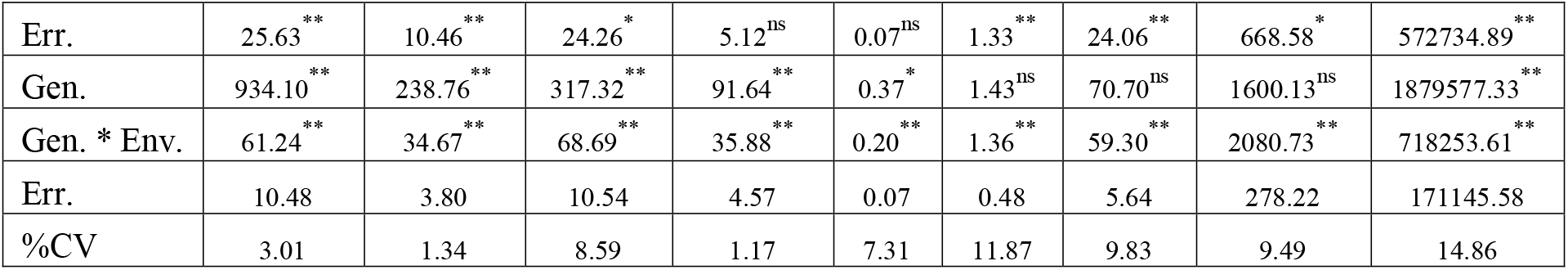
Combined ANOVA for 21 oilseed rape genotypes in the five tropical environments of Iran.

Additionally, evaluation of heritability showed phenological traits, including days to start flowering and days to end flowering had the highest heritability, which emphasizes their importance in oilseed rape selection and breeding (Table 4). Days to flowering trait has been previously shown the highest heritability with a value of 73.12%, while lower value (30.15%) for grain yield has been reported (Amiri-Oghan *et al.*, 2012), consistent with our results. Likewise, Naazar et al.,(Ali *et al.*, 2003) observed the highest and the lowest values of heritability for phenological and yield components traits. Phenological traits are the principal factors in oilseed rape breeding specially for the tropical regions because of high temperature at the late season that could lead to detriment in pod maturity and cause yield reduction. Yield improvement through manipulating of flowering time has been reported as one of the main strategies (Thurling & Kaveeta, 1992).Therefore, while phenological traits shows the highest heritability, it could be considered in the selection of early flowering and early maturity genotypes to escape from the late season heat stress.

**Table 4:** Evaluated heritability for quantitative traits of 21 oilseed rape genotypes in the five tropical environments of Iran.

|  | DSF | DEF | FP | DM | TSW | NSB | NGP | NPP | Y |
| --- | --- | --- | --- | --- | --- | --- | --- | --- | --- |
| Var G | 58.19 | 13.61 | 16.58 | 3.72 | 0.01 | 0.00 | 0.76 | 0<sup>*</sup> | 77421.58 |
| VarE | 0.70 | 0.25 | 0.70 | 0.30 | 0.00 | 0.03 | 0.38 | 18.55 | 11409.71 |
| Var G*E | 16.92 | 10.29 | 19.38 | 10.44 | 0.04 | 0.29 | 17.89 | 600.84 | 182369.34 |
| Var Pho | 75.81 | 24.15 | 36.66 | 14.46 | 0.06 | 0.33 | 19.02 | 587.35 | 271200.63 |
| %Hb | 76.76 | 56.33 | 45.21 | 25.71 | 18.23 | 1.42 | 4.00 | nd | 28.55 |

\*: negative estimated value of genotype variation is considered as zero.

Correlation analysis using the Person method indicated a significant relationship among some quantitative traits (Table 5). Days to flowering and maturity showed a positive correlation with one-thousand seed weight. On the other hand, days to flowering and maturity showed a negative correlation with the number of pods per plant. Therefore, delay in flowering and maturity lead the oilseed rape genotype to increase seed weight and decrease pod number.

**Table 5:** Correlation analysis for quantitative traits of 21 oilseed rape genotypes in the five tropical environments of Iran.

| Variables | DSF | DEF | FP | DM | TSW | NSB | NGP | NPP | Y |
| --- | --- | --- | --- | --- | --- | --- | --- | --- | --- |
| DSF | **1** |  |  |  |  |  |  |  |  |
| DEF | **0.885** | **1** |  |  |  |  |  |  |  |
| FP | **-0.712** | **-0.302** | **1** |  |  |  |  |  |  |
| DM | **0.888** | **0.891** | **-0.472** | **1** |  |  |  |  |  |
| TSW | **0.518** | **0.443** | **-0.391** | **0.487** | **1** |  |  |  |  |
| NSB | 0.058 | 0.061 | -0.028 | -0.044 | 0.004 | **1** |  |  |  |
| NGP | 0.029 | -0.063 | **-0.155** | 0.046 | 0.012 | 0.016 | **1** |  |  |
| NPP | **-0.149** | **-0.151** | 0.076 | **-0.305** | **0.145** | 0.109 | **-0.178** | **1** |  |
| Y | -0.077 | -0.023 | **0.122** | **-0.182** | -0.085 | **0.293** | 0.016 | **0.162** | **1** |
| Values in bold are different from 0 with a significance level alpha=0.05 |  |  |  |  |  |  |  |  |  |

The number of grain per pod showed a negative correlation with the number of pods per plant and flowering period. Thus, by increasing the number of pods in the plant, decreasing the grain number of the pod would be expected. Additionally, by extension of the flowering period, the number of grains per pod would be increased.

It has been also observed that flowering period length relates positively with a yield. A negative correlation was also observed between days to maturity and yield. The number of sub-branches and also the number of pods per plant also showed positive correlation with yield. Therefore, by increasing the flowering period, the number of sub-branches and the number of pods per plant, the oilseed rape yield would be increased. Our correlation results between yield and yield components coincided precisely with the previous published report in oilseed rape genotypes, which has been reported a significant positive correlation among yield and the number of pods per plant and the number of sub-branches but not a significant correlation with the number of grains per pod and one thousand seed weight(Baradaran *et al.*, 2007).

In order to investigate the direct and indirect effects of quantitative traits on yield, path analysis has been studied in the current study. Results showed that the days to maturity had the most negative direct effect on yield and the days to start flowering, while the number of sub-branches had the most positive direct effect on yield. One thousand seed weight and the number of pods per plant affected positively on yield indirectly through days to start flowering and the number of sub-branches, respectively (Figure 1).

**Figure 1:**
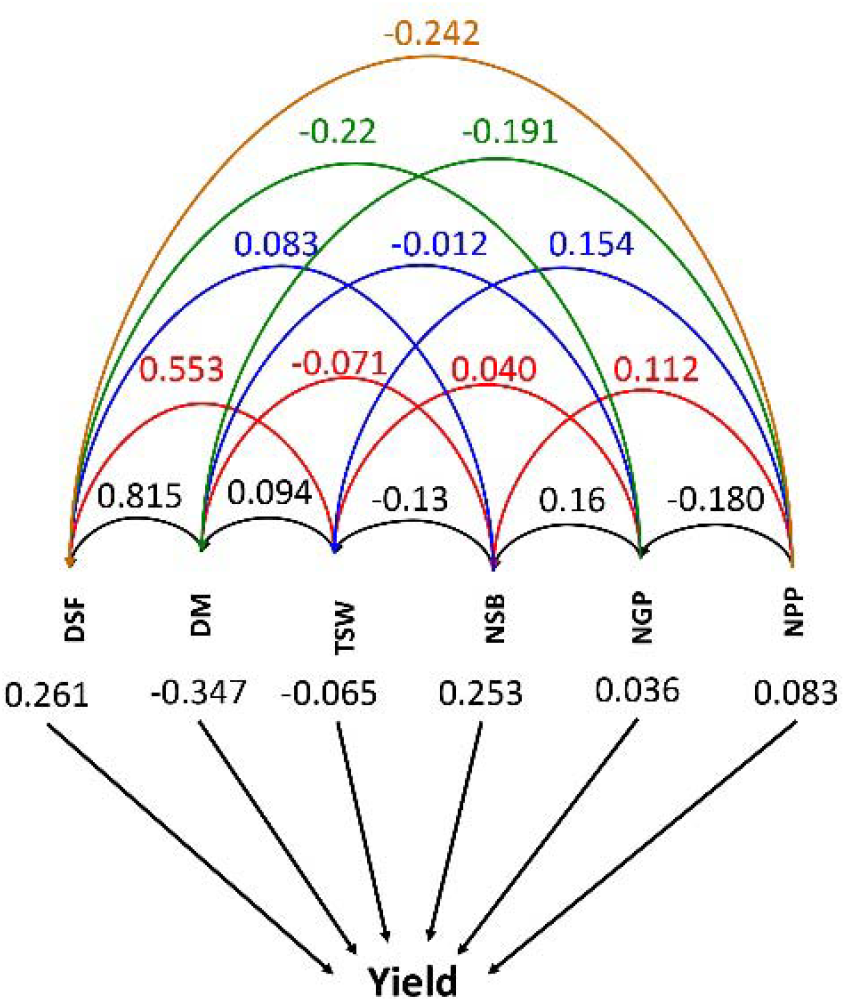
Path analysis for quantitative traits of 21 oilseed rape genotypes in the five tropical environments of Iran.

Canonical correlation analysis has been conducted to investigate the relationship between phonological traits and yield components (Figure 2). This analysis confirmed correlation and path analysis outputs. As shown in figure 2, the yield is correlated positively with phenological traits. Likewise, the number of sub-branches and the number of pods per plant were positively correlated with phenological traits. Therefore, analyzing Person correlation, canonical correlation and path analysis revealed phenological traits, especially days to maturity, are the main limiting factor in increasing yield in the five test locations in the tropical regions of Iran.

**Figure 2:**
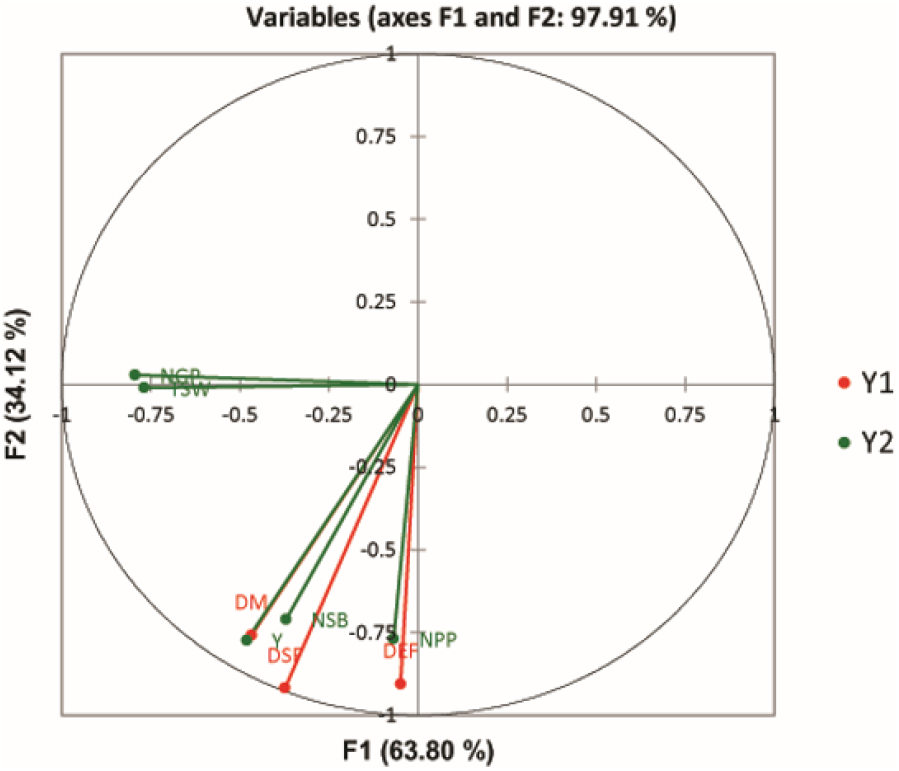
Canonical correlation for quantitative traits of 21 oilseed rape genotypes in the five experimental locations of Iran.

In the current study, the principal component analysis was performed according to evaluated quantitative traits for 21 oilseed rape genotypes. Results showed that the two first components covered 68.07% of all data variations. Therefore, a two-dimensional bi-plot was drawn based on the first two components (figure 3). 50.90 % of the variation was covered by the first component, which showed a strong positive correlation with days to start flowering, days to end flowering, days to maturity, number of sub-branches, number of pods per plant and yield. In this regard, it was named as a component of yield and phenological traits. The placement of yield and phenological traits together in the first component showed the close relationship of phenological traits on yield. Previous reports were also reported that the number of pods per plant and yield belonged to the first component (Baradaran *et al.*, 2007; Moradi *et al.*, 2017).The second component covered 17.17 % of the variation, which was positively correlated with one thousand seed weight and the number of grains per pod. This classification was consistent with canonical correlation results (Figure 2). According to bi-plot analysis, 12 of 21 genotypes were correlated with the first two components and separated, which had adequate yield in the five experimental locations. These genotypes were included G1-5, G12, G14-17, G19 and G21(Figure 3).

**Figure 3:**
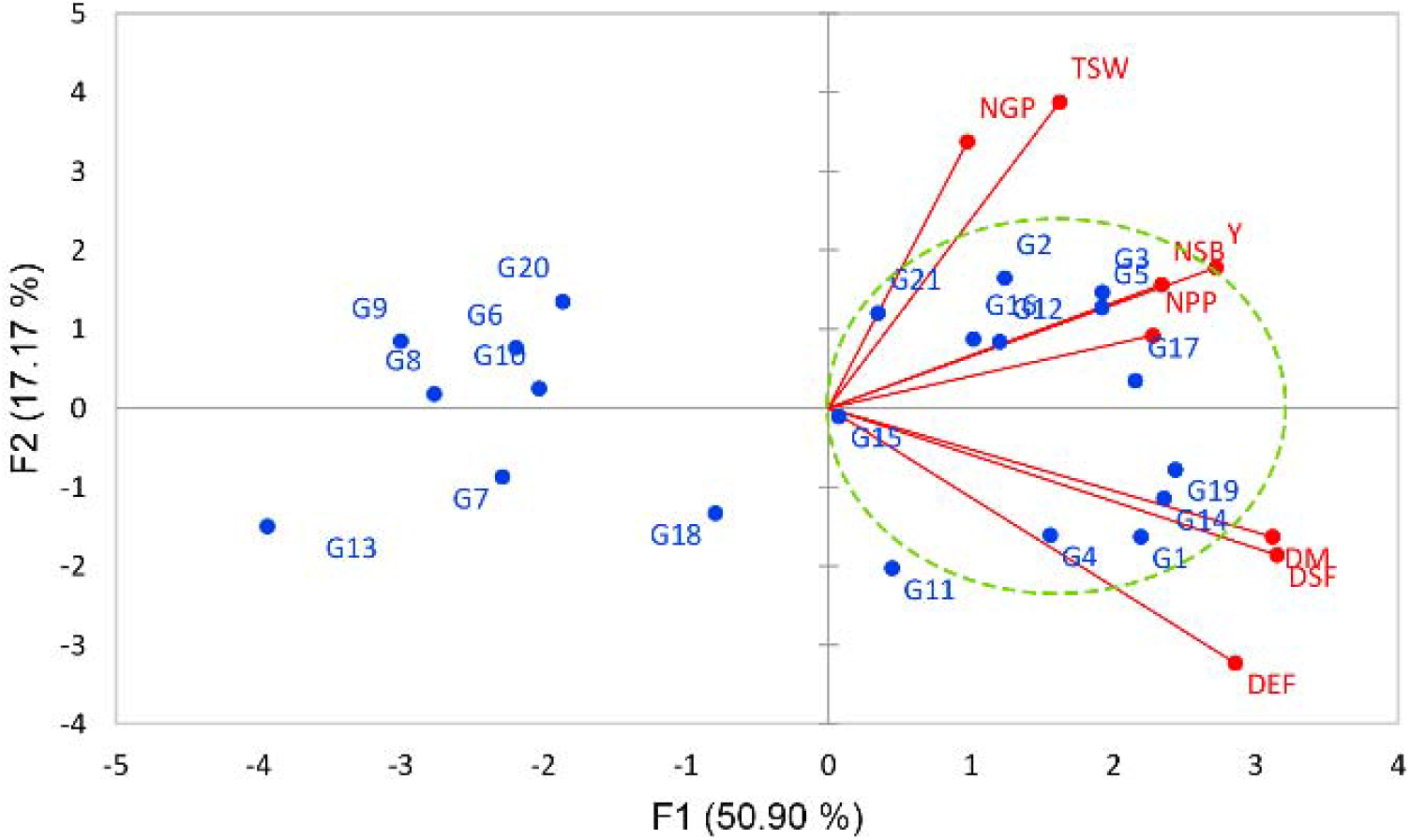
distribution of 21 oilseed rape genotypes based on two principal components and vectors of quantitative traits.

Cluster analysis has been conducted in the current study to classifying 21 oilseed rape genotypes using the WARD method. Cluster analysis categorized genotypes into three main groups. The first, second and third classes were included 8, 9 and 4 genotypes (Figure 4-a).To compare different classes for their quantitative traits, all traits values were converted to Z scores and then compared (Figure4-b). The first class showed the lowest values of days to start flowering, days to end flowering, days to maturity, one thousand seed weight, number of sub-branches, number of pods per plant and yield but the highest value of the flowering period. The second and third class showed the moderate values of days to start flowering, days to end flowering, days to maturity, but unlike the third class the number of grains per pod, the number of pods per plant and yield were high in the second class. Therefore, three classes, including early maturity genotypes with low yield (class 1), moderate maturity genotypes with high yield (class 2) and moderate yield and maturity date (class 3), were observed using cluster analysis. Most of the first class genotypes also showed no correlation with the first two components in bi-plot analysis, unlike the second class genotypes which showed the highest correlation with the two first components. Therefore, the second class genotypes included G2, G3, G5, G12, G14, G15, G17, G19 and G21 (check) were considered as promising genotypes with high yield and adaptability in the tropical regions in the current study.

**Figure 4:**
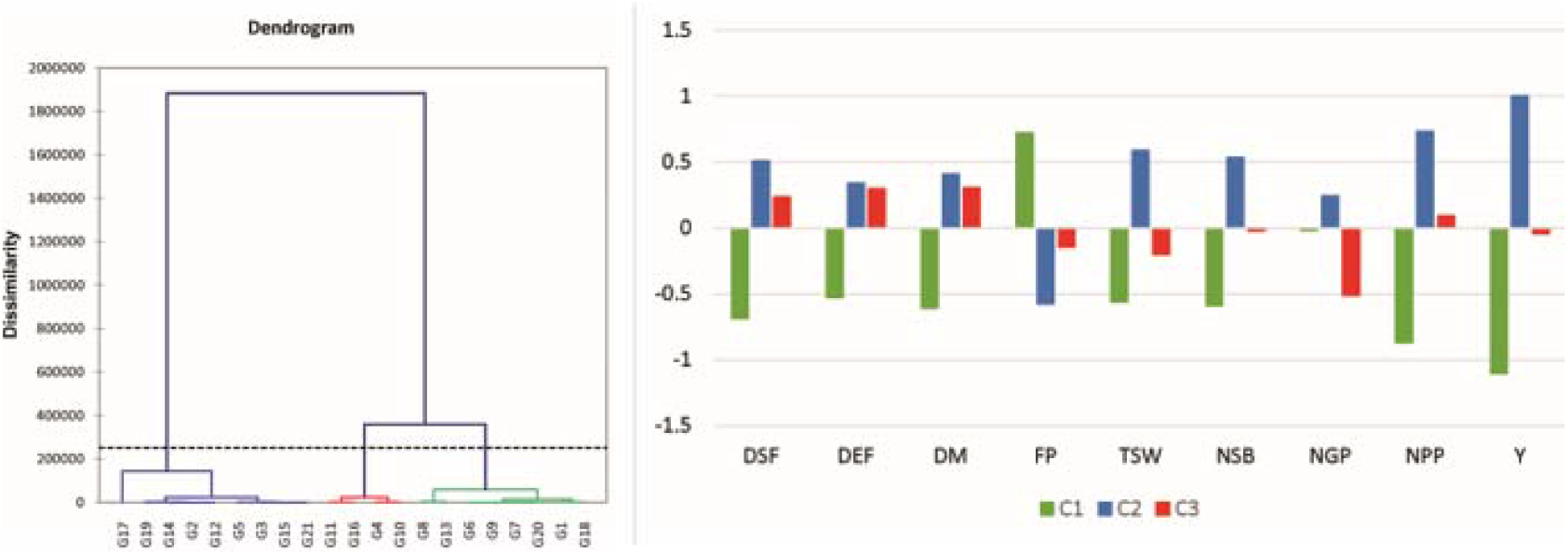
Cluster analysis for quantitative traits of 21 oilseed rape genotypes in the five tropical environments of Iran. The left figure showed the cluster analysis using the WARD method and the right figure showed a comparison of three classes of cluster analysis for quantitative traits of 21 oilseed rape genotype.

## Conclusions

In the current study, 20 genotypes and a check variety were investigated for their yield and adaptability. Person correlation, canonical correlation and path analysis revealed that flowering and maturity dates are the essential factors in the tropical regions of Iran, with the highest effect on yield. On the other hand, high heritability was observed for flowering and maturity dates. Therefore, oilseed rape breeding based on phenological traits would lead us to achieving high yield genotypes with the feature of phenological adaptability. Finally, using bi-plot and cluster analysis, eight promising genotypes were detected in the current study with high yield and adaptability in the tropical regions of Iran.

## Acknowledgements

The authors gratefully acknowledge Seed and Plant Institute Improvement (SPII) for the financial support that made possible the research with number 03-03-0304-279-981210 conducted there.

